# Genetic population subdivision of Blue Swimming Crab (*Portunus pelagicus*) across Indonesia inferred from mitochondrial DNA: implication to sustainable fishery

**DOI:** 10.1101/2020.10.07.329508

**Authors:** Hawis Madduppa, Rina Martaulina, Zairion, Resha Mukti Renjani, Mujizat Kawaroe, Nurlita Putri Anggraini, Beginer Subhan, Indri Verawati

## Abstract

The blue swimming crab (BSC), *Portunus pelagicus* (Linnaeus 1758), inhabits coastal areas of Southeast and East Asia, and is one of high fisheries commodity with export value for Indonesia and global market demand increasing annually. However, the data of genetic diversity and their spatial connectivity of populations in Indonesia are not yet known, which is important to inform unit stock management and sustainable fisheries. This study aimed to determine genetic diversity and differentiation of blue swimming crab across Indonesia populations under different Fishery Management Area, and their spatial genetic connectivity, as well as implications for sustainable fishery. A total of 297 individuals were amplified using cytochrome oxidase I mitochondrial DNA. This study shown highest value of haplotype and nucleotide diversity in the eastern part of Indonesia, where exploitation is relatively low. Significant genetic differentiation between populations (*F*_ST_ = 0.954; *p* < 0.001) and the fishery management regions (*F*_ST_ = 0.964; *p* < 0.001) were revealed. Low spatial connectivity was observed between populations in a distance of at least more than 60 kilometers. This study suggests that BSC populations in Indonesia likely have several unit stock, and preferably different fisheries management plan and action across the region thoroughly and simultaneously is effective for management and their sustainable conservation.

## Introduction

The blue swimming crab (*Portunus pelagicus* Linnaeus, 1758) inhabits coastal water of Southeast and East Asia (Lai *et al*. 2010). The spatial connectivity of blue swimming crab is influenced by their life cycle and other environmental parameters. Juvenile and adult of *P. pelagicus* inhabit benthic coastal environments for both estuaries and nearshore and females migrate into high salinity water for spawning, but they seem not return into the estuaries or nearshore after spawning (Zairion *et al*. 2014), except larval stage (Potter *et al*. 1983; Kangas 2000; Kembaren *et al*. 2018). Juvenile of *P. pelagicus* couldn’t find at the nearshore where water salinity less than 10 PSU (Kurnia *et al*. 2014), and this species would migrate from estuary to marine water as a reaction to lowered salinity (Meagher 1971; Potter *et al*. 1983, 1991). The blue swimming crab has life cycle with five larval stages, while larval duration might for 26–45 days, and then the crab phase (Kangas 2000). Since this species has moderately long planktonic larval stages and might has high mobility during the crab phase, it could be occure a high gene flow level within and between populations (Edgar 1990).

The Indonesian blue swimming crab fishery has been developed rapidly since 1990s and became an important source of coastal communities’ income. The crab meat has been exported approximately 20,000mt per annum over the last decade, primarily to USA markets. This product is demanding to ecolabel certification and traceability system now (Madduppa et al. 2016). However, according to the government and industrial meat production, the average size of blue swimming crabs landed has been declining since 2008. The catch per trip is also fluctuated; eventhough the unit and dimention of fishing gears have increased. This trend is also occurred in region where the blue swimming crab fisheries developed earlier, and most of blue swimming crab fishery tends to be overfishing. It’s seemed that the Indonesian blue swimming crab fishery trends are threatening the resources and economic sustainability. Therefore, Indonesia as one of the important supplier’s countries of raw materials crab canning industry, which currently is the one commodity that is very important for more than 180 thousand woman pickers in miniplant and about 90 thousand fishermen in Indonesia, making it necessary rescue efforts crab population from extinction.

The genetic diversity reveals gene flow between and within population, and therefore could use to determine the healthy of population (Santos *et al*. 2010), and determining stability and resilience of population (Ferguson *et al*. 1995). Due to excessive fishing and environmental changes, many stocks have been depleted and some species might become endangered (e.g., Musick *et al*. 2000; Hutchings and Reynolds 2004; Dankel *et al*. 2008), and would be influencing the changes in population genetic diversity (e.g., O’Brien 1994; Heino and Godø 2002; Reusch *et al*. 2005). The decreasing of genetic diversity might occure in long time before the actual effect is visible, which will affect on long-term of increased in genetic drift and then the loss of variability and adaptation ability (Hauser *et al*. 2002; Spielman *et al*. 2004). The genetic structure information of population could effectively lead the fishery management in sutainable point of view (Nishida *et al*. 1998; Jefri *et al*. 2015; Toha *et al*. 2016).

The small sizes wild-caught proportion of *P. pelagicus* has been increasing in last decade in Indonesia and suggests that this species is over-exploitation. Dependence of production from capture results can reduce the number of small crab population. When the rate of increase in fishing effort is not comparable with the growth of the fish resources, fish stocks will be reduced and the resulting decline in catches. This condition is known as biological overfishing (Sparre and Vanema 1998). Critical behavior of the crab is the development life cycle that occurs in some places. In the larval phase and spawning phase is in the open sea crab (off-shore), while the juvenile phase until the adults were in coastal waters (near-shore) (Kangas 2000). However, there have been lack information on genetic diversity and population subdivisions of *P. pelagicus* in Indonesia. Knowlegde and identification result of reproductively isolated and/or genetically differentiated populations within a species are importance for restocking programs (Carvalho and Hauser 1994; Conover *et al*. 2006), and for the reconstruction of an appropriate management (i.e. the harvest strategy and harvest control rules) in this species (Bryars and Adams 1999). Despite its high value, the population genetic structure of *P. pelagicus* in Indonesia has not been well studied unlike countries such as Malaysia, Australia and Thailand (Yap *et al*. 2002; Klinbunga *et al*. 2007; Chai *et al*. 2017). One of study concerning morphometric character and mitochondrial 16S rRNA sequence of *P. pelagicus* has conducted in Burru water, South Sulawesi as small part of Indonesian waters (Hidayani *et al*. 2015). Therefore, this study aimed to investigate the diversity and structure of genetic within and between populations of blue swimming crab (*P. pelagicus*) across Indonesia, and discuss the implications to blue swimming crab fishery sustainability.

## Materials and Methods

### Tissue collection

The tissue sampling was conducted at fish landing sites around the harbor in each population during several expeditions in 2015-2017. The tissue sample was collected by cutting a piece of small crab legs and stored in a 2 mL microtube containing 96% ethanol with a label contain identity of the sample. A total of 297 tissue samples was collected from eleven populations representing six of Fishery Management Regions in Indonesia (Table 1).

**Table 1.** Number of samples (n), number of haplotype (*H*_n_), haplotype diversity (*H*_d_), and nucleotide diversity (π) from eleven populations representing six of fishery management area (FMA) across Indonesia, showing code for site shown in Figure 1

| Code | FMA | Site | n | Hn | Hd | $\pi$ |
| --- | --- | --- | --- | --- | --- | --- |
| BAT | 571 | Batu Bara | 26 | 8 | 0.526 | 0.025 |
| 2 | 711 | Bangka Belitung | 22 | 6 | 0.745 | 0.026 |
| 3 | 712 | Pulau Lancang | 49 | 6 | 0.543 | 0.012 |
| 4 | 712 | Rembang | 47 | 5 | 0.723 | 0.016 |
| 5 | 712 | Pamekasan | 60 | 12 | 0.542 | 0.015 |
| 6 | 713 | Ekas, Lombok | 22 | 5 | 0.468 | 0.018 |
| 7 | 713 | Maros | 18 | 6 | 0.621 | 0.020 |
| 8 | 714 | Bombana | 14 | 4 | 0.396 | 0.008 |
| 9 | 714 | Pamandati | 12 | 2 | 0.167 | 0.003 |
| 10 | 714 | Kendari | 15 | 6 | 0.571 | 0.120 |
| 11 | 715 | Halmahera | 12 | 6 | 0.758 | 0.034 |

### DNA extraction, amplification and sequencing

Extraction of genomic DNA for each sample was conducted by using extraction kit (Geneaid kit). A fragment of mitochondrial Cytochrome Oxidase subunit-I gene (COI) was amplified using the following primer set: CO1F 5’ AGA AGT GTA TAT TTT AAT TC -3’ and CO1R 5’-ATG TAG AAT ATC GAT AG-3’ (Folmer et al. 1994). Polymerase Chain Reaction (PCR) was conducted in 25 µl reaction volume containing 1-4 µl templates DNA, 2.5 µl of 10x PCR buffer (Applied Biosystems), 2.5 μL dNTP (8 mM), 2 µl MgCl_2_ (25 mM), 0.125 µl AmplyTaq Red™ (Applied Biosystems), 1.25 µl of each primer (10 mM), 1 µl 1x BSA, and 13.5 µl ddH_2_O. PCR conditions were: initial denaturation at 94°C for 15 s, followed by 40 cycles of denaturation at 95°C for 30s, annealing at 40°C for 30s, and extension at 72°C for 45s. The final extension step was conducted at 72°C for 10 min. The quality of PCR products was assessed by agarose gel electrophoresis and ethidium bromide staining and visualized using UV transilluminator. All good PCR products were sent to Sanger sequencing facility.

### Data Analysis

Mt-DNA sequences were aligned and edited using Mega 6 (Tamura *et al*. 2013). GENETYX program was used to calculate genetic distance (D) among individual in intra population and inter populations. Based on Kimura 2-parameter model and 1,000 replication of bootstrap was contructed a Neighbour-Joining (NJ) tree in Mega 6 (Tamura *et al*. 2013). Analyzing of a number of the haplotype (H) and haplotype diversity (Hd) by using DnaSP 5.10 (Nei 1987; Rozaz *et al*. 2003), while nucleotide diversity (π) followed the method of Lynch and Creasef (1990). Subsequently, fixation index (*F*_ST_) is used to assess population differentiation (Excoffier *et al*. 1992) and determined by Arlequin 3.5 (Excoffier and Lischer 2010). In order to investigate the phylogenetic relationship among haplotype, a minimum spanning tree was constructed in Network 4.6.1 (http://www.fluxusengineering.com). Then, Analysis Molecular Variance (AMOVA) was applied to assess genetic differentiation (*F*_ST_) among and within between populations based on sites and fishery management area (FMA) of blue swimming crab (*Portunus pelagicus*) across Indonesia (Excoffier *et al*. 1992; Roewer *et al*. 1996). The *F*_ST_ will show whether between sites and fishery management area (FMA) is genetically different (different unit stock) or panmixing.

## Results

### Genetic diversity

A total of 40 haplotype observed in all populations, where highest was found in Pamekasan and the lowest in Pamandati (Table 1). The highest value of haplotype and nucleotide diversity was observed in Halmahera (0.758) and Kendari (0.120), respectively. Pamandati was observed as the lowest for both haplotype (0.167) and nucleotide (0.003) diversity (Table 1).

### Population genetic structure

The reconstruction of phylogenetic trees *Portunus pelagicus* markers CO1 using Neighbour Joining (NJ) Kimura 2 parameter models with 1000 bootstrap values of eleven populations across Indonesia using 297 sequences showing five clades (Figure 2). The five clades as follows: (1) Java Island population (FMA 712), (2) Sulawesi dan Lombok (713-714), (3) North Sumatera (571), (4) Halmahera (715), and (5) Bangka Belitung (711).

**Figure 1.**
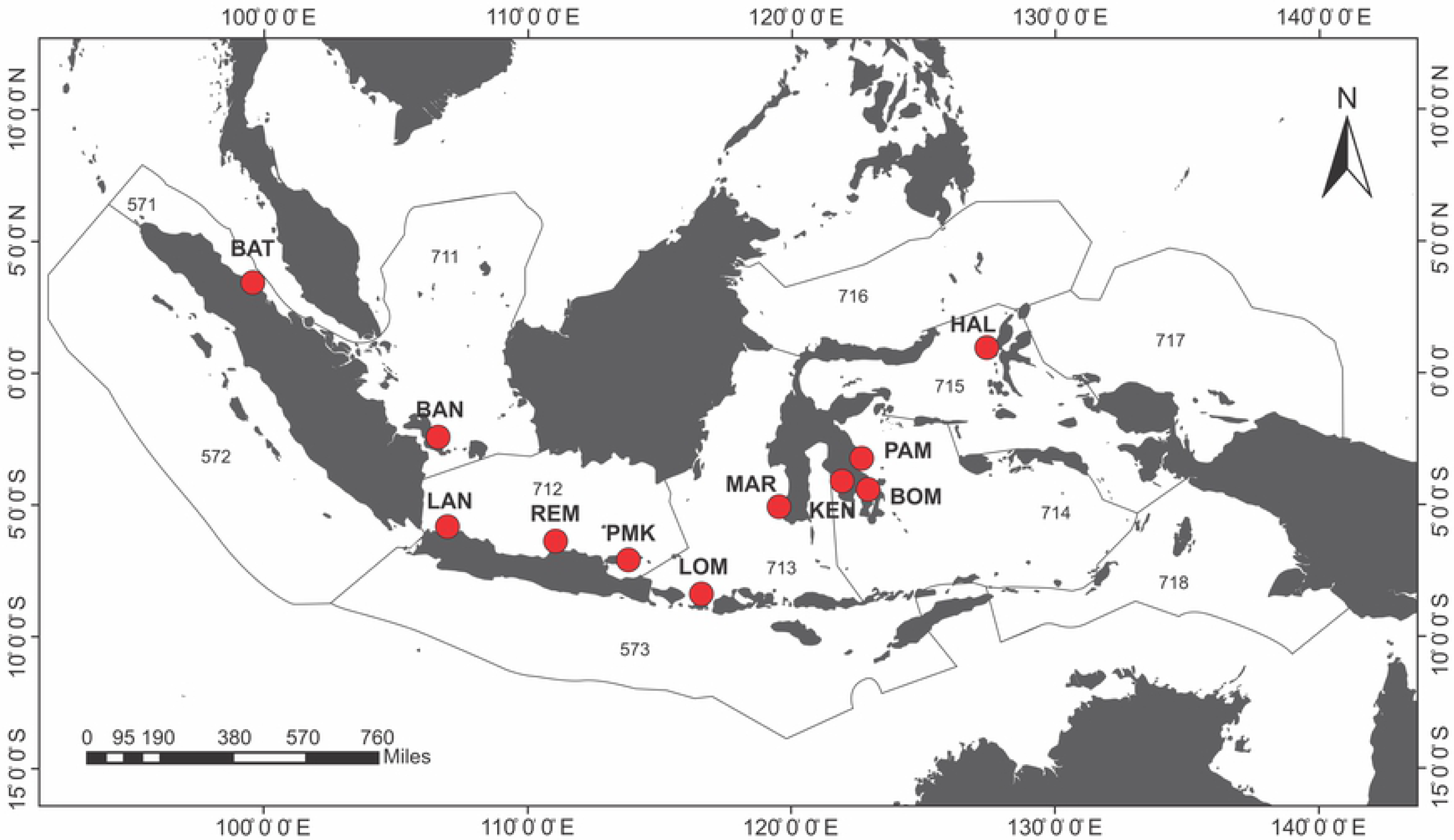
Tissue sample collection sites of blue swimming crab (*Portunus pelagicus*) at eleven populations (BAT: Batubara, BAN: Bangka Belitung, LAN: Pulau Lancang, REM: Rembang, PMK: Pamekasan, LOM: Ekas Lombok, MAR: Maros, BOM: Bombana, PAM: Pamandati, KEN: Kendari, HAL: Halmahera), representing six of fishery management areas (FMA) in Indonesia; FMA 571; 711; 712; 713; 714 and FMA 715.

**Figure 2.**
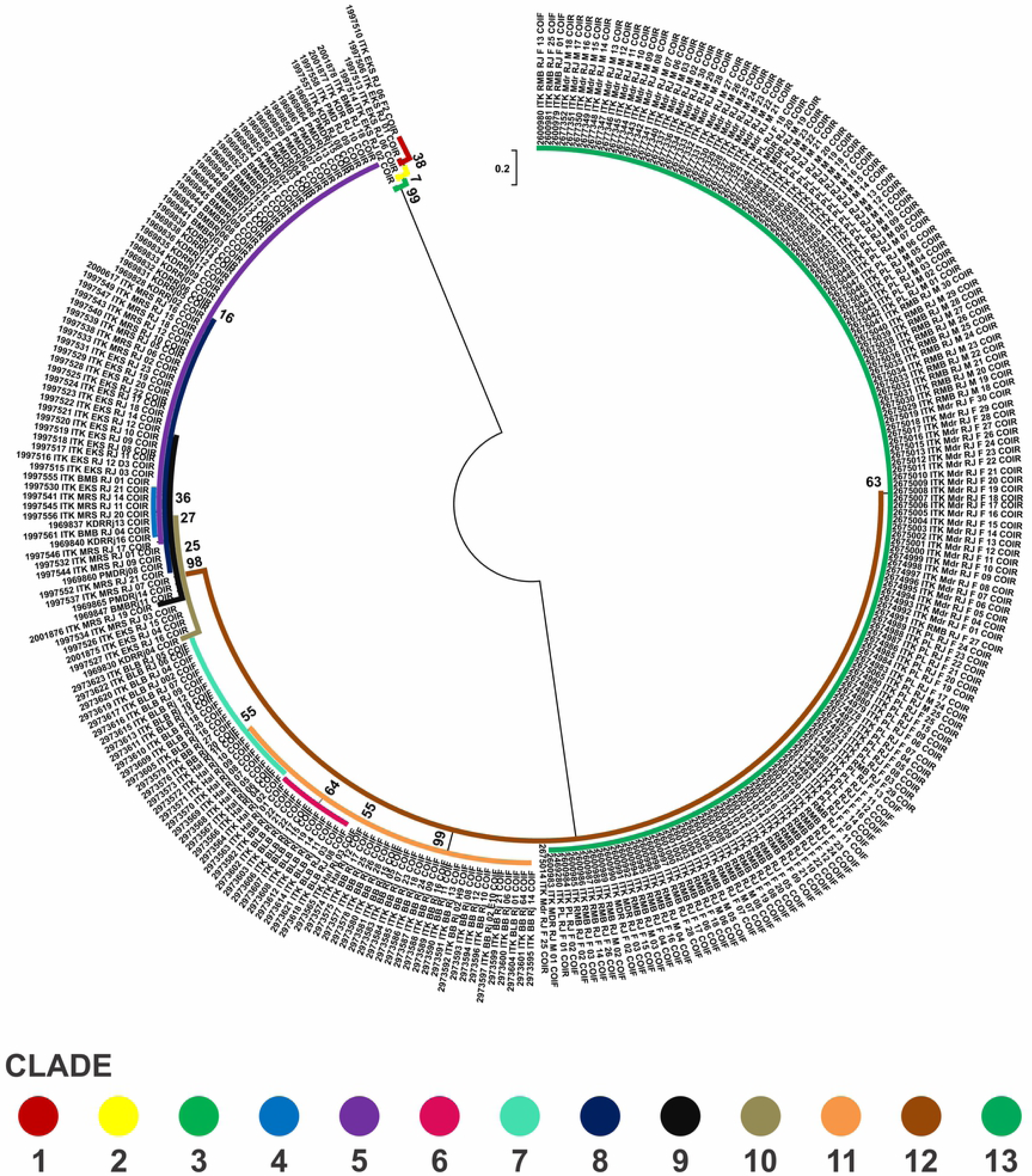
Reconstruction of phylogenetic trees *Portunus pelagicus* markers CO1 using Neighbour Joining (NJ) Kimura 2 parameter models with 1000 bootstrap values of eleven populations across Indonesia using 297 sequences showing five clades: (1) Java Sea population (FMA 712), (2) Sulawesi dan Lombok (FMA 713-714), (3) North Sumatera (FMA 571), (4) Bangka Belitung (FMA 711), and (5) Halmahera (FMA 715) from 13 hyplotypes.

The highest difference in genetic structure (*F*_ST_) among populations was shown between Bombana and Rembang of 0.9917, followed by populations between Kendari and Lancang (0.9860), and the closest populations between Kendari and Bombana (−0.0001) (Table 2). The results of the analysis of AMOVA (Analysis of Molecular Variance) showed strong genetic subdivision among eleven populations (*F*_ST_ = 0.954, p<0.001) and Fishery Management Area (*F*_ST_ = 0.964, p<0.001) (Table 3). The haplotype network of blue swimming crab (*P. pelagicus*) using model TCS Network from a total of 297 sequences of eleven populations across Indonesia showed that haplotype was distributed across Indonesia, but a significant difference FMA. The haplotype network, distribution and composition of blue swimming crab (*Portunus pelagicus*) using model TCS Network from a total of 297 sequences of eleven populations across sampling sites in Indonesia show that each site dominated by one haplotype (Figure 3 and Figure 4). Figure 5 and Figure 6 show that FMA 712 was separated differently to western Indonesia areas (FMA 571 and 711), and eastern Indonesia areas (FMA 713, 714, and 715).

**Table 2.** The genetic differentiation shown by Fixation Index (*F*_ST_) among sites of *Portunus pelagicus* populations across study sites in Indonesia

| Sites | Ekas | Maros | Lancang | Rembang | Pamekasan | Kendari | Bombana | Pamandati | Halmahera | Batu Bara |
| --- | --- | --- | --- | --- | --- | --- | --- | --- | --- | --- |
| Maros | -0.0041 |  |  |  |  |  |  |  |  |  |
| Lancang | 0.9373 | 0.9345 |  |  |  |  |  |  |  |  |
| Rembang | 0.9353 | 0.9323 | 0.1494 |  |  |  |  |  |  |  |
| Pamekasan | 0.9411 | 0.9388 | -0.0119 | 0.1390 |  |  |  |  |  |  |
| Kendari | 0.8236 | 0.8044 | 0.9860 | 0.9853 | 0.9860 |  |  |  |  |  |
| Bombana | 0.8417 | 0.8238 | 0.9922 | 0.9917 | 0.9915 | -0.0001 |  |  |  |  |
| Pamandati | 0.8409 | 0.8229 | 0.9920 | 0.9915 | 0.9913 | 0.0075 | -0.0775 |  |  |  |
| Halmahera | 0.8812 | 0.8682 | 0.9749 | 0.9736 | 0.9748 | 0.9662 | 0.9815 | 0.9809 |  |  |
| Batu Bara | 0.9084 | 0.9022 | 0.9731 | 0.9721 | 0.9731 | 0.9731 | 0.9822 | 0.9812 | 0.4585 |  |
| Bangka | 0.9056 | 0.8984 | 0.97708 | 0.9760 | 0.9768 | 0.9748 | 0.9856 | 0.9852 | 0.0714 | 0.6424 |

**Table 3.** Fixation index (*F*_ST_) value of blue swimming crab (*Portunus pelagicus*) in Fisheries Management Area (FMA) of Indonesia

| FMA | 571 | 711 | 712 | 713 | 714 |
| --- | --- | --- | --- | --- | --- |
| 711 | 0.6364 |  |  |  |  |
| 712 | 0.9773 | 0.9793 |  |  |  |
| 713 | 0.8825 | 0.8808 | 0.9479 |  |  |
| 714 | 0.9135 | 0.9117 | 0.9639 | 0.7029 |  |
| 715 | 0.4464 | 0.7140 | 0.9785 | 0.8602 | 0.8962 |

**Table 4.** Result of Analysis of Molecular Variance (AMOVA) and Fixation Index (*F*_ST_) based on sites and fishery management area (FMA) of blue swimming crab (*Portunus pelagicus*) across Indonesia

| Source of variation | d.f. | Variation (%) | $F_{ST}$ | Significance ($p$) |
| --- | --- | --- | --- | --- |
| Sites |  |  |  |  |
| Among populations | 10 | 95.44 | 0.954 | <0.001 |
| Within populations | 286 | 4.56 |  |  |
| FMA |  |  |  |  |
| Among populations | 5 | 96.36 | 0.964 | <0.001 |
| Within populations | 291 | 3.64 |  |  |

**Figure 3.**
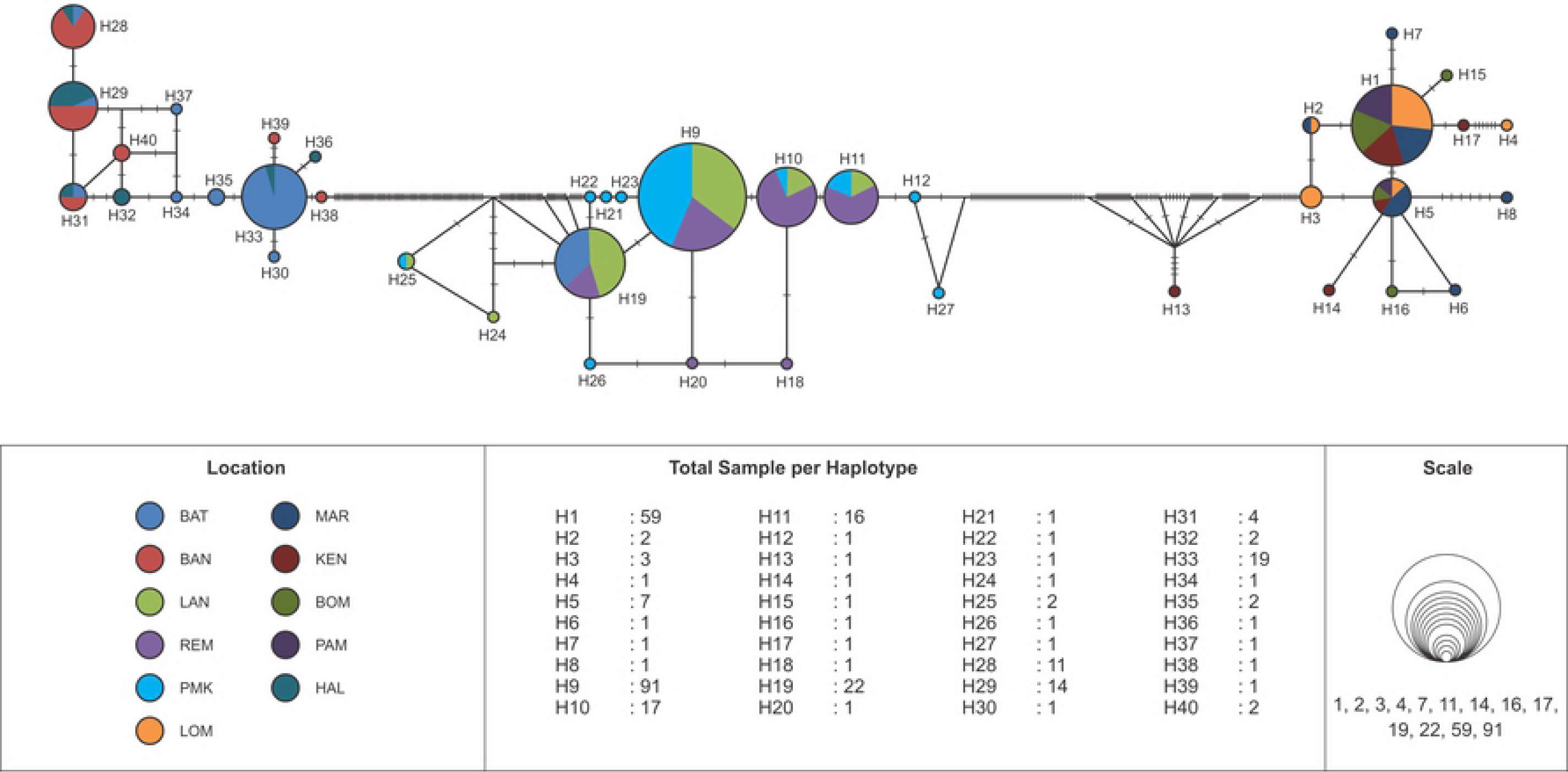
The haplotype network of blue swimming crab (*Portunus pelagicus*) using model TCS Network from a total of 297 sequences of eleven populations across sampling sites in Indonesia

**Figure 4.**
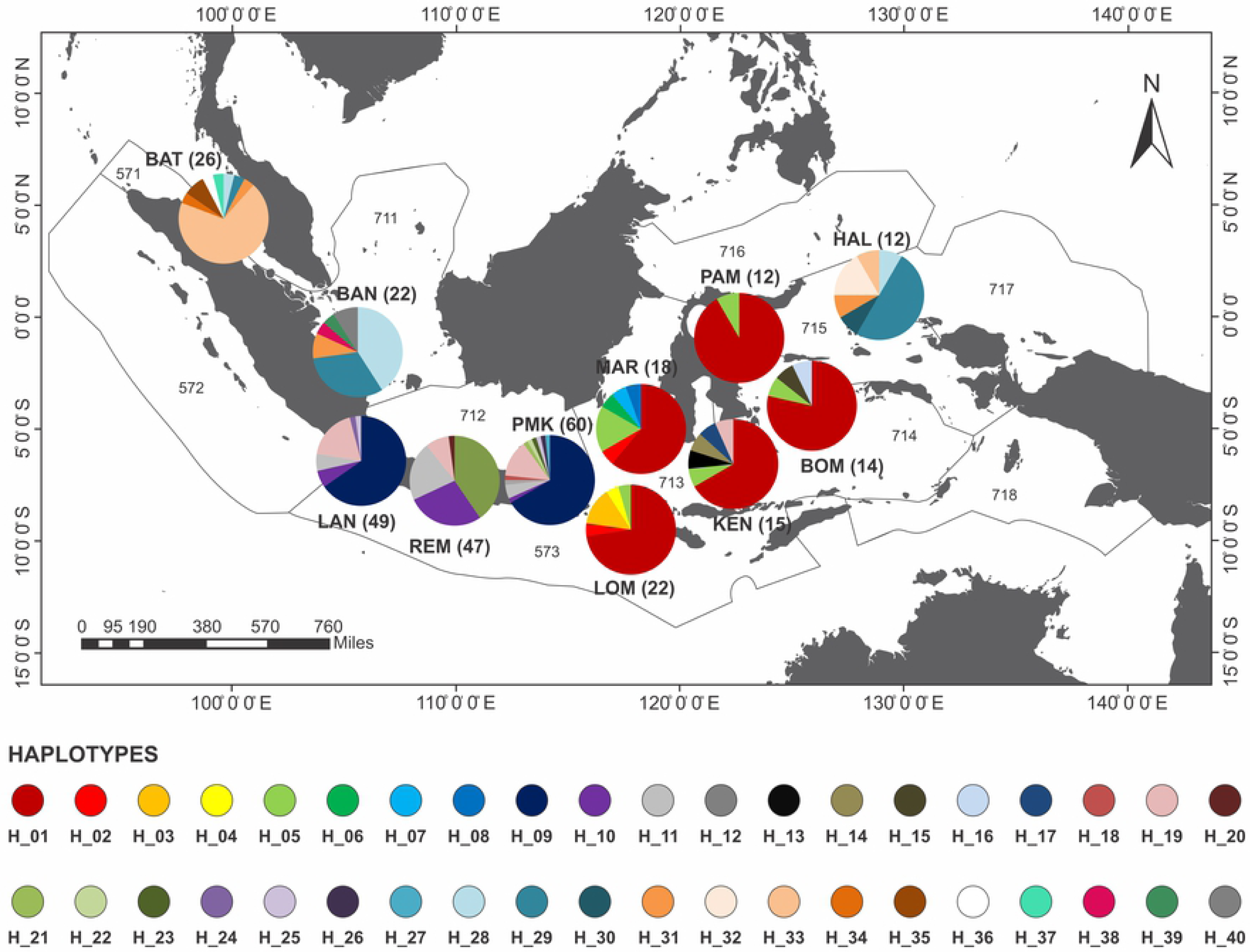
The haplotype distribution and composition of blue swimming crab (*Portunus pelagicus*) using model TCS Network from a total of 297 sequences of eleven populations across sampling sites in Indonesia. In bracket is number of sample per site.

**Figure 5.**
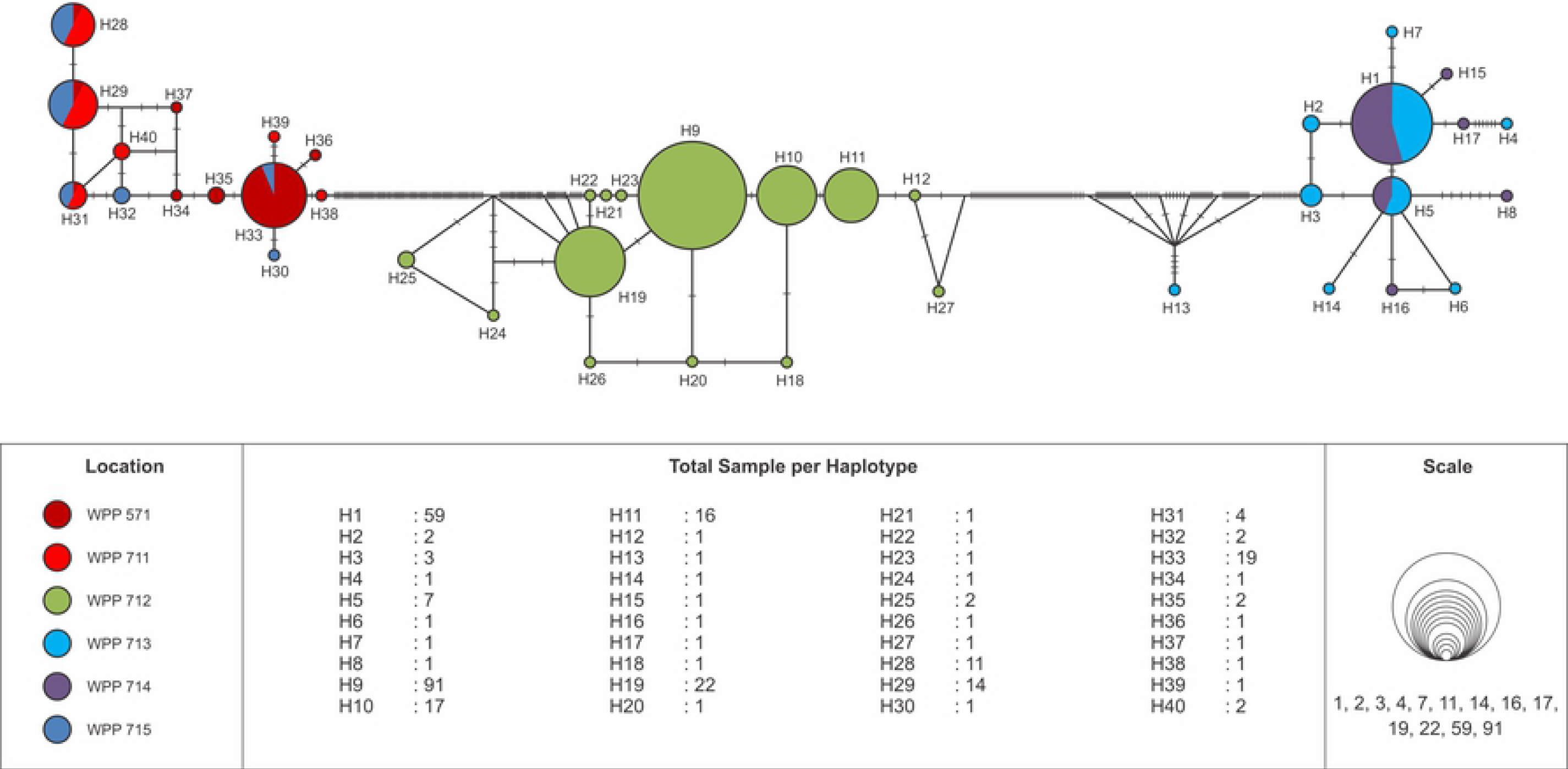
The haplotype network of blue swimming crab (*Portunus pelagicus*) using model TCS Network from a total of 297 sequences of eleven populations across Indonesia within fisheries management region.

**Figure 6.**
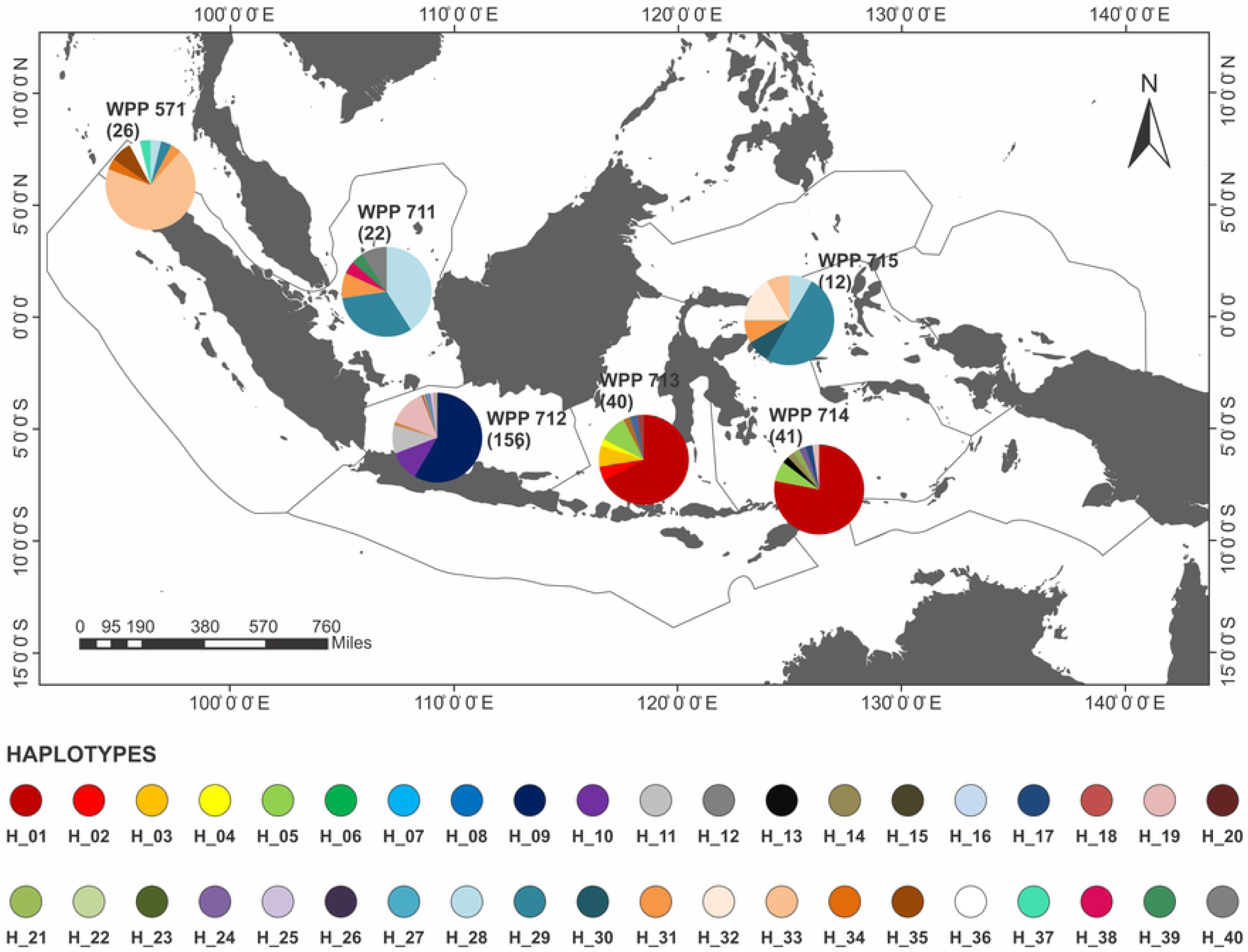
The haplotype distribution and composition of blue swimming crab (*Portunus pelagicus*) using model TCS Network from a total of 297 sequences of eleven populations across Indonesia within fisheries management region. In bracket is number of sample per region.

## Discussion

### Genetic diversity

The value of the population genetic diversity is affected by the number of haplotypes (Hn) that exist. This study shown highest value of haplotype and nucleotide diversity in the eastern part of Indonesia (Halmahera and Kendari). A large number of haplotypes is directly proportional to the value of genetic diversity. The low genetic diversity of *P. pelagicus* population is also observed in at the site of the Gulf of Saint Vincent, Spencer Gulf, and coastal areas of Western Australia (Sezmiş 2004), because the number of haplotypes on each individual is dominated by a single haplotype. Meanwhile, on the east coast and the port of the western Australian waters found high genetic diversity, because of the large number of different haplotypes. Similar result observed in Thailand that showed a high haplotype diversity and low nucleotide diversity (Ren *et al*. 2018). A total of 40 haplotype observed in all populations, where the highest was found in Pamekasan and the lowest in Pamandati. Total haplotype has diverse type of impact on the genetic diversity in a population. The more diverse types of composite haplotype population level of genetic diversity, genetic in the population will be higher and vice versa (Smith and Chesser 1981). Genetic diversity is affected by the transfer of genetic material between populations of different locations (Slarkin 1987). Genetic diversity increases when there is a genetic input from other populations, called genetic migration. Great Migration would lead to interbreeding and mixing of genes between different populations, in order to obtain variations in different genes (Lande and Barrowdough 1987). The presence of a wide variety of genes of individuals in the population would be increasing the population ability in response to the changes of environmental condition. High genetic diversity in the population of fish may protect from environmental interferences (Hartl and Jones 1998). Taylor and Aarsen (1988) explains that species with good adaptability will produce variations in phenotype and genotype both in response to particular environmental conditions so as to increase the ability of individuals to survive and proliferate.

The amount of genetic diversity observed in study sites may be due to fishing pressure regularly and crab are in a stage of self - environmental adaptation, with a population that is still available in nature, but are vulnerable to extinction. This case seemed to be happen in area of Southeast Sulawesi where the fishing pressure are high, and also some areas in the north coast of Java except Madura where still be found a high genetic diversity. The loss of genetic diversity will reduce the population ability to adapt to environmental changes (Frankham 1996). Genetic diversity has a potential impact directly or indirectly on the population, and might follow the impact to the community and ecosystem (Hughes *et al*. 2008). Gen a living being is a fundamental reproductive unit in the formation of the individual, so the decline in genetic diversity will decrease the population for sustainable success. Moreover, population size should be estimated in determining the appropriate amount of catches (Madduppa *et al*. 2014).

### Population genetic structure

The reconstruction of phylogenetic trees for *Portunus pelagicus* on this study showing distinct five clades, of each Fisheries Management Area (FMA), except in Sulawesi and Lombok (713-714). Sezmiş (2004) produced similar results, i.e., the formation of two major clade of *P. armatus* (previously known as *P. pelagicus*), each clade consisting of individual mixes of different populations in Australian waters. Phylogenetic interpreted as a model for representing approximately relationship ancestor organism, molecular sequences or both (Brinkman and Leipe 2001). This shows that although individual crab apart in different populations, but a third of this population is closely related genetically or derived from a common ancestor. In addition, these results indicate that the crab migration patterns in the third phase of life of the population are still in the same location. Geographical distance is still within the scope of the population of the province strengthened the case of the results obtained. Branching tree pattern matrix formed by the distance between pairs that can describe the population genetic fusion that occurs in the group (Weiss 1995).

A high value of *F*_ST_ was indicating strong subdivision and low gene flow between populations. The current study is similar to some of previous studies, which have revealed sub structure on the *P. pelagicus* populations. In Australia, some studies showed Australian waters are devided into four subpopulations based on different markers (i.e. allozyme polymorphism, COI and microsatellite). The study of Bryars and Adams (1999) using allozyme showed subpopulations between populations (such as West Coast, Spencer Gulf, and Gulf St. Vincent in South Australia and Darwin-Gove in the Northern Territory), and within each subpopulation among the sites. Other population genetic structure of *P. pelagicus* was investigated from 16 different assemblages throughout the geographic range of this species in Australian waters (Sezmiş 2004). This investigation used six microsatellite loci and a 342 bp fragment of cytochrome oxidase subunit I (COI), and found significant genetic heterogeneity.

The current study is contradictory as previously assumed by Yap *et al*. (2002) that the *P. pelagicus* who has an extended planktonic larval stages would potentially has high larval dispersal and might occure of extensive gene flow between conspecific samples within geographic mesoscale that is from ten to hundred of kilometres. In contrast with the observed result in China, population genetic structure is limited occure with a high level of gene flow along the distribution areas of this species in China (Ren *et al*. 2018). While in Malaysia, microsatellites analyses indicated low levels of genetic differentiation among the *P. pelagicus* populations (Chai *et al*. 2017). This result also supports that *P. pelagicus* are capable of moving substantial distances, with one recorded as traveling 20 km in one day in Moreton Bay, Queensland (Smith and Sumpton 1989 Sumpton *et al*. 1994). Other observation concerning *P. pelagicus* is also regarded as a potential species vagile since adults are able to travel in geographic mesoscale daily (Kangas 2000), which showed by the current study a mixed population within a region. It seems that within a mesoscale regions, blue swimming crab population is tend to be mixed as also showed by the current study (e.g. within Java Sea region), but will be different genetically in great scale.

### Implications to fishery management

The blue swimming crab has high productivity and rapid growth rates (Ernawati *et al*. 2017), thus, depleted blue swimming crab stocks could recover quickly by restoring and maintaining breeding sized crabs in the stock. The strong subdivision and low gene flow between populations as shown by a high value of *F*_ST_, suggest that each fisheries management area (FMA) in Indonesia should be treated differently by developing fishery management plan and strategic action at each management region. The biological characteristics of blue swimming crab, the coherent organized nature of the industry, and its reliance on sustainability conscious export markets makes the blue swimming crab fishery strategic for starting the process and inclusive collaborative management model for the coastal fisheries (Ghofar et al. 2018). Information on the genetic diversity of this crab can be further used as a database on mapping potential area of Indonesia blue swimming crab. Crab management in each population needs to be done thoroughly and simultaneously. It is required in the implications for sustainable crab fishery, because it can determine an effective strategy that is continues maintained. This study should provide insight into population genetic structure of *P. pelagicus* in Indonesia, including conservation genetics, and utilization management of this species in sustainable basis.

## Acknowledgement

The research was funded by the Indonesian Blue Swimming Crab Association (APRI) and the Laboratory of Marine Biodiversity and Biosystematics Department of Marine Sciences and Technology Faculty of Fisheries and Marine Science, Bogor Agricultural University.

